# Multistage Machine Learning Reveals Circadian Gene Programs and Supports a Retina–Choroid Axis in Myopia Development

**DOI:** 10.64898/2026.04.02.716020

**Authors:** Akarapon Watcharapalakorn, Teera Poyomtip, Patarakorn Tawonkasiwattanakun, Putri Krishna Kumara Dewi, Thotsapol Thomrongsuwannakij, Tanakamol Mahawan

**Author notes:** Correspondence: Tanakamol Mahawan.

## Abstract

**Purpose:** To determine whether circadian timing defines critical molecular windows in myopia development and to assess the transferability of circadian gene programs across ocular tissues, disease stages, and species.

**Methods:** Publicly available retinal and choroidal RNA-seq datasets from chick models of form-deprivation myopia were analyzed using unsupervised transcriptomic profiling and multistage machine-learning classification. Circadian windows were defined based on Zeitgeber time, and samples were grouped accordingly for downstream analyses. Classification model robustness was evaluated through cross-tissue and cross-stage validation and further assessed using external validation in an independent dataset. Functional translation to humans was examined using ortholog-based Gene Ontology enrichment analysis to identify conserved biological processes and higher-order regulatory pathways.

**Results:** A circadian critical window at ZT8–ZT12 exhibited the strongest transcriptional divergence during both myopia onset and progression. Gene signatures derived from this window generalized across retina and choroid and remained predictive across disease stages, supporting coordinated molecular regulation between ocular tissues. External validation confirmed the reproducibility of these signatures despite differences in experimental design and gene coverage. Functional mapping revealed that conserved molecular components in chicks are reorganized into more complex neuroendocrine and regulatory networks in humans, indicating cross-species conservation with increased functional complexity.

**Conclusions:** Circadian timing strongly shapes myopia-related gene expression and underlies coordinated retina–choroid signaling. These findings highlight circadian biology as a key factor of refractive development and suggest that time-dependent mechanisms may influence myopia susceptibility, progression, and response to treatment.

## 1. Introduction

Myopia (nearsightedness) is a major global public health concern, with projections suggesting that nearly half of the world’s population will be affected by 2050 [1]. In addition to the burden of refractive error, myopia—particularly high myopia—is associated with an increased risk of sight-threatening complications, including retinal detachment, myopic maculopathy, glaucoma, and choroidal neovascularization [2, 3]. Despite extensive research, the biological mechanisms underlying abnormal ocular elongation remain incompletely understood.

Refractive development is a visually guided process in which the retina detects optical defocus and initiates signaling cascades that propagate through the choroid to the sclera, ultimately regulating eye growth[4]. This retina–choroid–sclera signaling axis integrates visual input, neuromodulation, metabolic regulation, and extracellular matrix remodeling [5]. Notably, pathological changes are often first observed in the retina, supporting its central role in initiating growth-related responses[6].

Circadian biology has increasingly been recognized as an important regulator of ocular growth[7]. In chicks and mice, manipulating circadian timing or light–dark cycles alter ocular growth. Similarly, in humans, reduced daylight exposure and irregular sleep patterns increase the risk of myopia onset, suggesting the involvement of daytime light and circadian entrainment in myopia development[1,2]. However, the molecular mechanisms linking circadian regulation to refractive development remain poorly defined.

The chick (*Gallus gallus domesticus*) is a well-established model for visually induced myopia[12]. Form deprivation myopia (FDM) can be reliably induced using a translucent diffuser, producing rapid axial elongation and retinal and choroidal changes that resemble human myopia. Importantly, chicks exhibit robust circadian rhythms in retinal physiology, making them suitable for investigating time-of-day– dependent mechanisms[13].

Recent transcriptomic studies have characterized diurnal gene expression in chick retina and choroid during myopia onset and progression, revealing pronounced time-of-day–dependent differences between occluded and control eyes [14, 15]. These findings suggest that circadian phase modulates the molecular response to visual deprivation. However, these datasets have not been systematically integrated, and it remains unclear whether specific circadian windows—such as ZT8 and ZT12—represent critical phases of myopia-related signaling.

Machine learning (ML) approaches have been increasingly used in myopia research to identify molecular biomarkers and model complex gene expression patterns associated with ocular growth [16, 17]. By capturing nonlinear and multivariate relationships in high-dimensional data, ML enables robust classification and cross-dataset validation of myopia-related molecular signatures, typically using differential expression and feature selection methods. While these approaches have identified candidate biomarkers, they are generally based on single datasets and do not incorporate temporal or circadian structure [18]. More recent models integrating clinical and metabolomic features have shown promising predictive performance, but they do not account for circadian dynamics[19]. Thus, a key gap remains in applying integrative, multi-stage ML frameworks to circadian transcriptomic data in myopia. In this study, we integrate publicly available retinal and choroidal RNA-seq datasets using a multistage ML framework to identify circadian-critical windows of gene expression in myopia. We further evaluate the transferability of these gene signatures across tissues, disease stages, and independent datasets, and assess their functional relevance through cross-species analysis. By linking circadian regulation to retina–choroid signaling, this work provides new insight into the temporal organization of ocular growth and its role in myopia development.

## 2. Materials and Methods

### 2.1 Data Collection

This study reanalyzed publicly available retinal and choroidal RNA-seq datasets from previously published myopia studies (Stone et al., 2024 [14, 15]), including **GSE227724** (myopia onset) and **GSE261232** (myopia progression). Both datasets were generated from Cornell K strain White Leghorn chicks subjected to form-deprivation myopia under 12 h light/12 h dark conditions. The onset dataset represents one light–dark cycle, while the progression dataset represents four consecutive cycles. Retina and choroid tissues were collected at six Zeitgeber time points (ZT0–ZT20, 4 h intervals; n = 6 per time point). Raw sequencing data and expression matrices were obtained from Gene Expression Omnibus (GEO; https://www.ncbi.nlm.nih.gov/geo/) and used for downstream analyses.

### 2.2 Circadian Critical Window Identification

We hypothesized that ZT8–ZT12 represents a circadian critical period based on previous studies showing that this mid-day interval contains the largest number of differentially expressed genes in both retina and choroid during experimental myopia. Importantly, those earlier analyses compared open versus occluded eyes, demonstrating that ocular growth signals are most transcriptionally dynamic around ZT8– ZT12.

Building on this biological rationale, the present study uses ZT8–ZT12 as the classification target for machine-learning modeling, with all other time points grouped as ZT_other as shown in Figure 1.This approach tests whether multistage ML can identify gene programs that characterize this hypothesized critical window, rather than defining the window from the current dataset itself.

**Figure 1.**
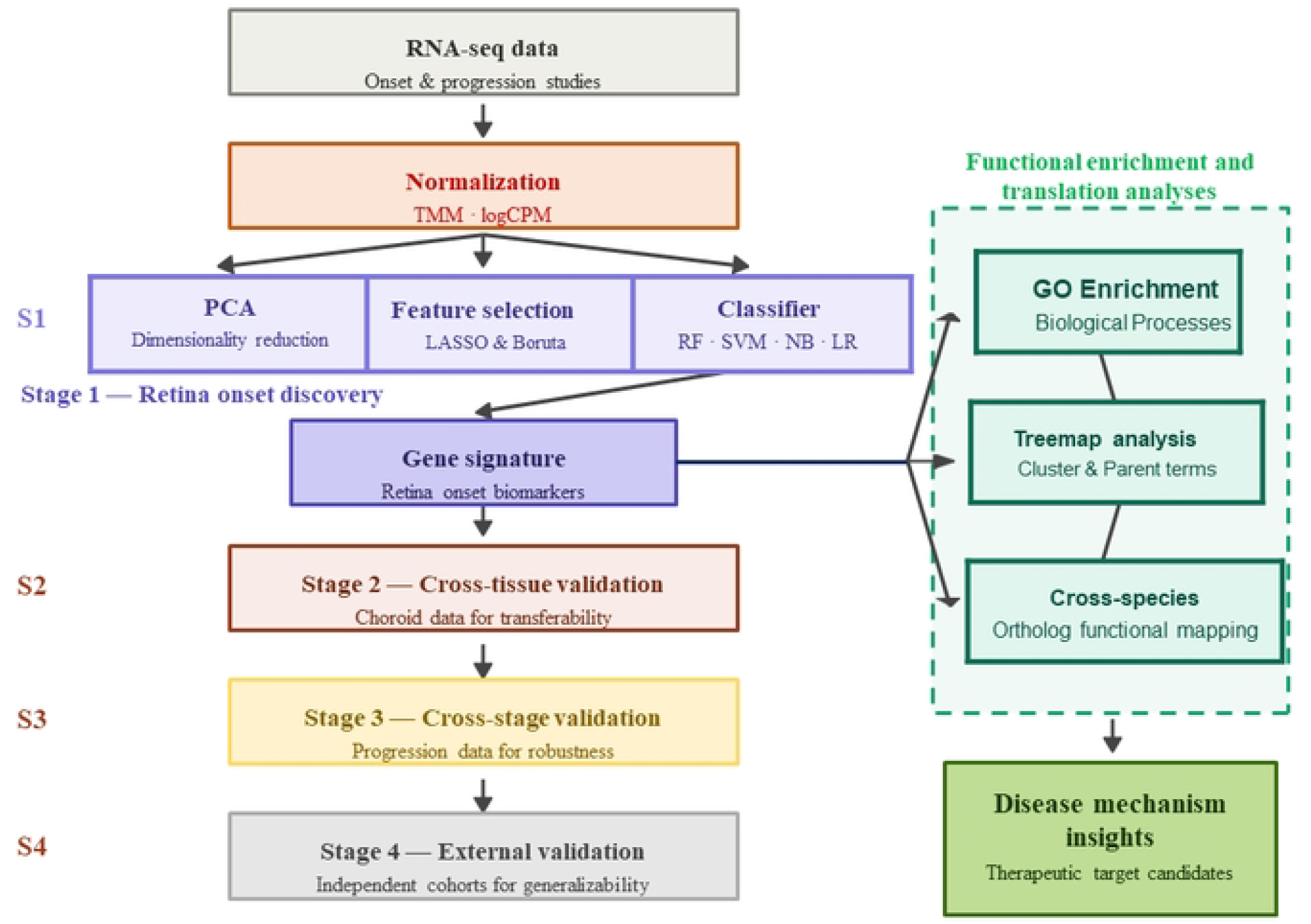

### 2.3 Machine Learning Analysis

The machine learning framework consisted of four stages: (i) feature discovery in the retina onset dataset, (ii) cross-tissue validation in the choroid onset dataset, (iii) cross-stage validation using independent progression datasets, and (iv) external validation using independent external datasets as demonstrated in Figure 1. All data analysis and modelling were analyzed by R programming language version 4.5.1 [20, 21].

#### 2.3.1 Stage 1: Primary Discovery Model (Retina—Onset Dataset)

RNA-seq count matrices and sample metadata were obtained from GSE227724. Raw counts were processed using the *edgeR* package (version 4.6.3) [22], with low-expression genes filtered using *filterByExpr* and library sizes normalized using the Trimmed Mean of M-values (TMM) method. Data were transformed to log2 counts per million (logCPM). Zeitgeber time (ZT) was encoded as a circular variable using sine and cosine transformations (sin_time = sin(2πZT/24), cos_time = cos(2πZT/24)) to capture circadian periodicity. Samples were categorized as ZT_812 or ZT_other, and sex was included as a covariate.

The dataset was partitioned into training (80%) and test (20%) sets using stratified sampling. Feature selection and model development were performed exclusively on the training set to prevent data leakage. Two feature selection methods were applied: LASSO logistic regression using the *glmnet* package (version 4.1.10) [23] and Boruta using the *Boruta* package (version 9.0.0) [24]. Feature selection was repeated 50 times with different random seeds, and only genes consistently selected in all iterations were retained. Three gene sets were evaluated: LASSO-selected, Boruta-selected, and overlapping genes.

Classification models were developed using Random Forest (RF), Support Vector Machine (SVM), Naïve Bayes (NB), and Logistic Regression (LR) implemented in the *caret* package (version 7.0.1) [25]. Model training was repeated 50 times using stratified resampling (80% training, 20% testing). Performance was evaluated using Accuracy, Precision, Recall, F1-score, area under the receiver operating characteristic curve (AUC), and Matthew’s correlation coefficient (MCC), with summary statistics and 95% confidence intervals calculated across iterations.

Gene signatures derived from the retina onset dataset were subsequently evaluated for cross-tissue and cross-stage transferability.

#### 2.3.2 Stage 2: Cross-Tissue Validation (Retina → Choroid)

To assess transferability across tissues, gene signatures derived from the retina onset dataset were applied to choroid samples from the same study. Models were trained using the same algorithms (RF, SVM, NB, LR) and evaluated using identical procedures, including 50 iterations of stratified resampling. Performance was assessed using the same metrics (Accuracy, Precision, Recall, F1-score, AUC, MCC).

#### 2.3.3 Stage 3: Cross-Stage Validation (Onset → Progression)

To evaluate temporal robustness, retina-derived gene signatures were applied to the progression dataset. Models were trained using the same algorithms and circadian classification (ZT_812 vs ZT_other). Classification was repeated 50 times using stratified resampling, and performance was assessed using the same evaluation metrics to determine whether signatures identified during onset remained predictive during progression.

#### 2.3.4 Stage 4: External Validation (Independent Dataset)

External validation was conducted using an independent retinal RNA-seq dataset from a myopia study (**GSE203604**; *n* = 42). Gene expression data were processed and normalized prior to analysis, and exploratory structure was assessed using unsupervised approaches. Predictive performance of the circadian gene signature was evaluated using nested leave-one-out cross-validation with regularized logistic regression and linear support vector machine models. Model performance was quantified by the AUC, with statistical significance assessed by permutation testing and confidence intervals estimated by bootstrap resampling. Temporal trends across experimental days were further examined using linear modeling.

### 2.4 Functional enrichment and translation analyses

To investigate the biological processes underlying myopia-associated gene signatures across species, Gene Ontology (GO) enrichment analysis was performed on Boruta-selected genes in chicken and their corresponding human orthologs using the *clusterProfiler* package (version 4.16.0) [26]. Significant GO terms (adjusted *p* ≤ 0.05) were summarized and reduced for redundancy using the *rrvgo* package (version 1.20.0) [27]., which groups related terms into representative parent GO categories.

Parent GO terms for chicken and human datasets were visualized using treemaps. To further characterize cross-species functional relationships, enrichment results were summarized at the level of parent GO biological processes and visualized using alluvial diagrams generated with the *alluvial* package (version 12.5.1) [28].

## 3. Results

### 3.1 Study Design and Sample Size of Data

The myopia onset and progression studies were conducted using identical circadian sampling schemes. For the onset study, retinal tissues were collected after one cycle of form deprivation on Day 1, while the progression study involved tissue collection after four cycles on Day 5. In both studies, samples were collected at six Zeitgeber time points (ZT0, ZT4, ZT8, ZT12, ZT16, and ZT20), with six biological replicates per time point, yielding a total of 144 samples per study as shown in **Table 1**.

**Table 1.** Experimental design, sampling scheme, and circadian classification.

| Study | Duration | Day | ZT Points | Rep/ZT | Total Samples | Class | n (%) |
| --- | --- | --- | --- | --- | --- | --- | --- |
| Onset | 1 cycle | 1 | ZT0, 4, 8, 12, 16, 20 | 6 | 144 | ZT_812 | 48 (33.3) |
|  |  |  |  |  |  | ZT_other | 96 (66.7) |
| Progression | 4 cycles | 5 | ZT0, 4, 8, 12, 16, 20 | 6 | 144 | ZT_812 | 48 (33.3) |
|  |  |  |  |  |  | ZT_other | 96 (66.7) |

**Table 2.** Performance of machine-learning classifiers for identifying the circadian critical window (ZT8/12) in the retina onset dataset with confidence interval 95%.

| Metric | RF | SVM | NB | LR |
| --- | --- | --- | --- | --- |
| **LASSO (n=24)** |  |  |  |  |
| Accuracy | 0.975 [0.931–1.000] | 0.949 [0.887–1.000] | 0.966 [0.915–1.000] | 0.889 [0.800–0.978] |
| Precision | 0.963 [0.910–1.000] | 0.899 [0.814–0.984] | 0.938 [0.870–1.000] | 0.809 [0.698–0.920] |
| Recall | 0.965 [0.913–1.000] | 0.965 [0.913–1.000] | 0.970 [0.922–1.000] | 0.880 [0.788–0.972] |
| F1-Score | 0.960 [0.905–1.000] | 0.923 [0.848–0.998] | 0.949 [0.886–1.000] | 0.830 [0.724–0.936] |
| AUC | 0.973 [0.927–1.000] | 0.954 [0.894–1.000] | 0.967 [0.917–1.000] | 0.887 [0.797–0.977] |
| MCC | 0.946 [0.882–1.000] | 0.895 [0.808–0.982] | 0.929 [0.857–1.000] | 0.764 [0.644–0.884] |
| **BORUTA (n=53)** |  |  |  |  |
| Accuracy | 0.986 [0.953–1.000] | 0.983 [0.947–1.000] | 0.985 [0.950–1.000] | 0.715 [0.587–0.843] |
| Precision | 0.964 [0.911–1.000] | 0.956 [0.898–1.000] | 0.964 [0.911–1.000] | 0.546 [0.405–0.687] |
| Recall | 1.000 [1.000–1.000] | 1.000 [1.000–1.000] | 0.995 [0.975–1.000] | 0.765 [0.645–0.885] |
| F1-Score | 0.980 [0.940–1.000] | 0.976 [0.932–1.000] | 0.977 [0.935–1.000] | 0.625 [0.488–0.762] |
| AUC | 0.990 [0.962–1.000] | 0.988 [0.957–1.000] | 0.988 [0.956–1.000] | 0.694 [0.564–0.824] |
| MCC | 0.972 [0.925–1.000] | 0.965 [0.914–1.000] | 0.968 [0.918–1.000] | 0.439 [0.299–0.579] |

Abbreviations: Random Forest (RF), Support Vector Machine (SVM), Naïve Bayes (NB), and Logistic Regression (LR)

**Table 3.** Cross-stage and cross-tissue validation performance of machine-learning classifiers.

| Metric | Cross-Stage Validation (Onset → Progression) |  |  |  | Cross-Tissue Validation (Onset: Retina → Choroid) |  |  |  |
| --- | --- | --- | --- | --- | --- | --- | --- | --- |
|  | RF | SVM | NB | LR | RF | SVM | NB | LR |
| Accuracy | 0.931 | 0.972 | 0.847 | 0.403 | 0.954 | 0.898 | 0.826 | 0.618 |
| Precision | 1.000 | 0.923 | 1.000 | 0.358 | 0.937 | 0.846 | 0.696 | 0.428 |
| Recall | 0.792 | 1.000 | 0.542 | 1.000 | 0.925 | 0.855 | 0.925 | 0.615 |
| F1 | 0.884 | 0.960 | 0.703 | 0.527 | 0.925 | 0.837 | 0.774 | 0.512 |
| AUC | 0.896 | 0.979 | 0.771 | 0.448 | 0.946 | 0.886 | 0.827 | 0.630 |
| MCC | 0.847 | 0.941 | 0.664 | 0.193 | 0.898 | 0.777 | 0.684 | 0.226 |

*Abbreviations: Random Forest (RF), Support Vector Machine (SVM), Naïve Bayes (NB), and Logistic Regression* *(LR)*

### 3.2 Circadian-Critical Windows of Gene Expression Divergence

Based on exploratory transcriptomic analyses, samples collected at ZT8 and ZT12 were grouped and defined as the circadian critical window (ZT_812), comprising 48 samples (33.3%) in each study. The remaining time points (ZT0, ZT4, ZT16, and ZT20) were classified as ZT_other, representing 96 samples (66.7%) as shown in Table 1, and were used as the comparison group for downstream machine-learning analyses. The detailed phenotype data including ZT class was found in **Supplementary Data1**.

The PCA revealed a distinct clustering of samples collected at ZT8 and ZT12, indicating shared transcriptomic characteristics that were separable from other time points as shown in Figure 2. This observation was further supported by preliminary differential expression analyses, which showed that ZT8 and ZT12 contained the highest number of differentially expressed genes in both retina and choroid tissues during myopia onset and progression.

**Figure 2.**
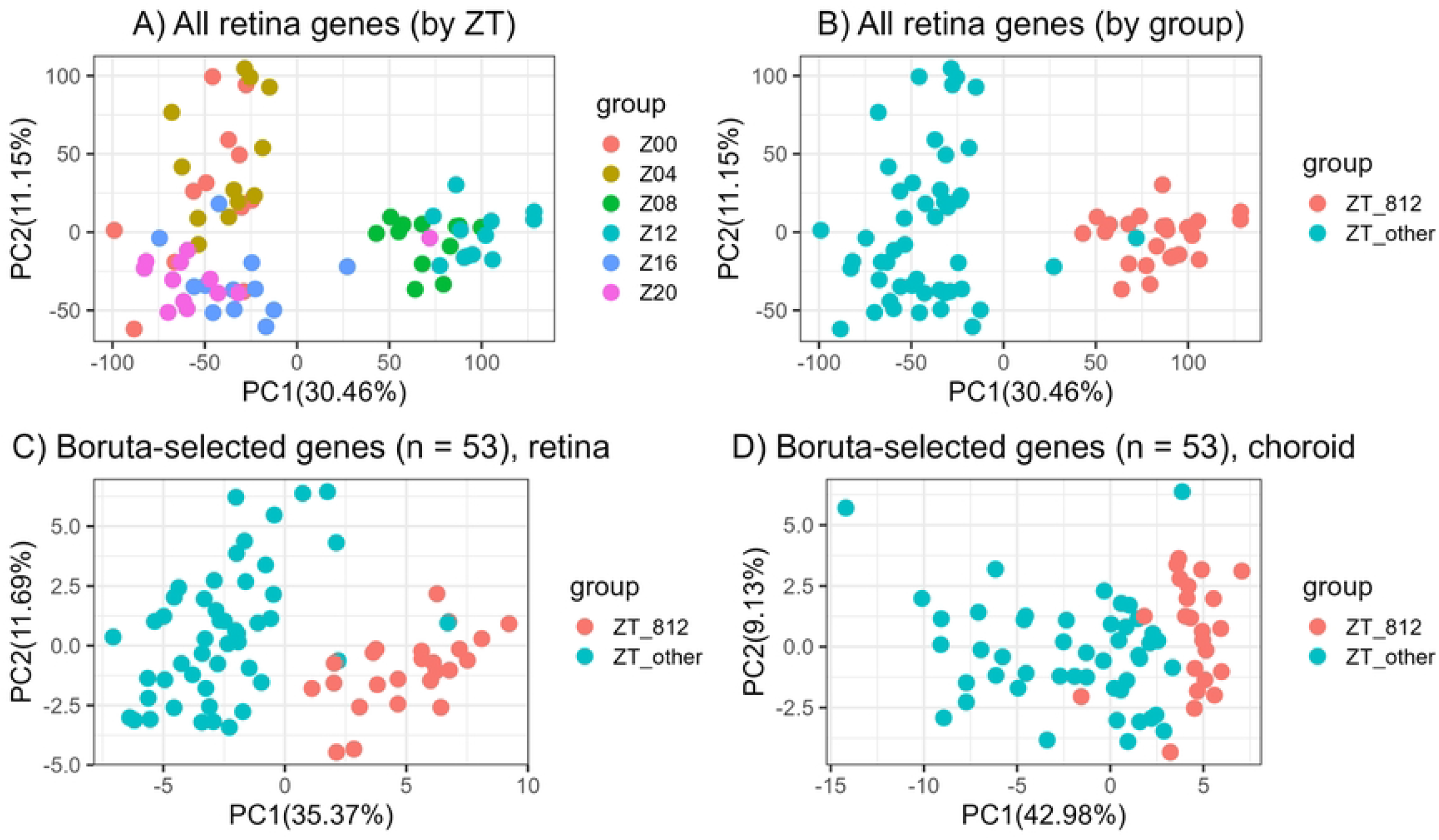

### 3.3 ML Classification Accurately Identifies Circadian-Critical Windows (Stage 1)

Feature selection was performed exclusively on the retina onset dataset to identify early molecular signals associated with myopia development. After 50 repeated runs, LASSO selected 24 genes, while Boruta identified 53 genes that were consistently selected in all iterations (50/50). These gene sets were subsequently used as input features for supervised machine learning classification. The 53 gene ID and their orthologs were listed in **Supplementary Data 2**.

Supervised models trained on the retina onset dataset effectively distinguished samples collected during the circadian critical window (ZT8/12) from other time points. Using the LASSO-selected gene set (n = 24), all classifiers showed high performance, with RF achieving the highest accuracy (0.975) and AUC (0.973), followed by NB and SVM. LR showed comparatively lower performance.

Using the Boruta-selected gene set (n = 53), performance further improved for RF, SVM, and NB, with RF achieving near-perfect discrimination (accuracy = 0.986; AUC = 0.990). In contrast, LR performance declined with the larger feature set, suggesting reduced robustness to higher-dimensional input. Overall, RF and Boruta selected genes demonstrated superior performance across feature sets, supporting the reliability of the identified circadian gene signatures. Therefore, we used Boruta selected genes to further validated in Stage2-4.

### 3.4 Cross-Dataset and Cross-Tissue Validation (Stage 2 and 3)

To assess the generalizability of the circadian gene signature, models trained on the retina onset dataset were evaluated in two independent validation settings: cross-stage validation using the progression dataset and cross-tissue validation using choroidal samples. In the cross-stage setting (onset → progression), SVM achieved the highest overall performance (accuracy = 0.972; AUC = 0.979), followed by RF, which also demonstrated strong discrimination (accuracy = 0.931; AUC = 0.896).

In the cross-tissue validation setting (onset retina → choroid), RF demonstrated the most robust performance across all metrics (accuracy = 0.954; AUC = 0.946), indicating strong transferability of the circadian gene program between retinal and choroidal tissues. Across both validation scenarios, RF and SVM consistently achieved higher MCC, reflecting balanced sensitivity and specificity and supporting the robustness of the identified circadian signature across disease stages and tissues.

### 3.5 External Validation on Independent Data (Stage 4)

Unsupervised analyses based on these genes showed consistent expression structure, with PCA indicating separation of samples across time points and experimental groups in Figure 3(b). Nested LOOCV analysis showed an AUC of 0.823 for logistic regression and 0.918 for the linear SVM as shown in Figure 3(a). Permutation testing indicated that the SVM model performed significantly better than chance (p = 0.015), whereas logistic regression did not (p = 0.68). Bootstrap confidence intervals were 0.782–0.997 for logistic regression and 0.841–0.998 for SVM. The composite signature score increased across time points from D1 to D6 particularly in PC1 as depicted in Figure 3(c-d).

**Figure 3.**
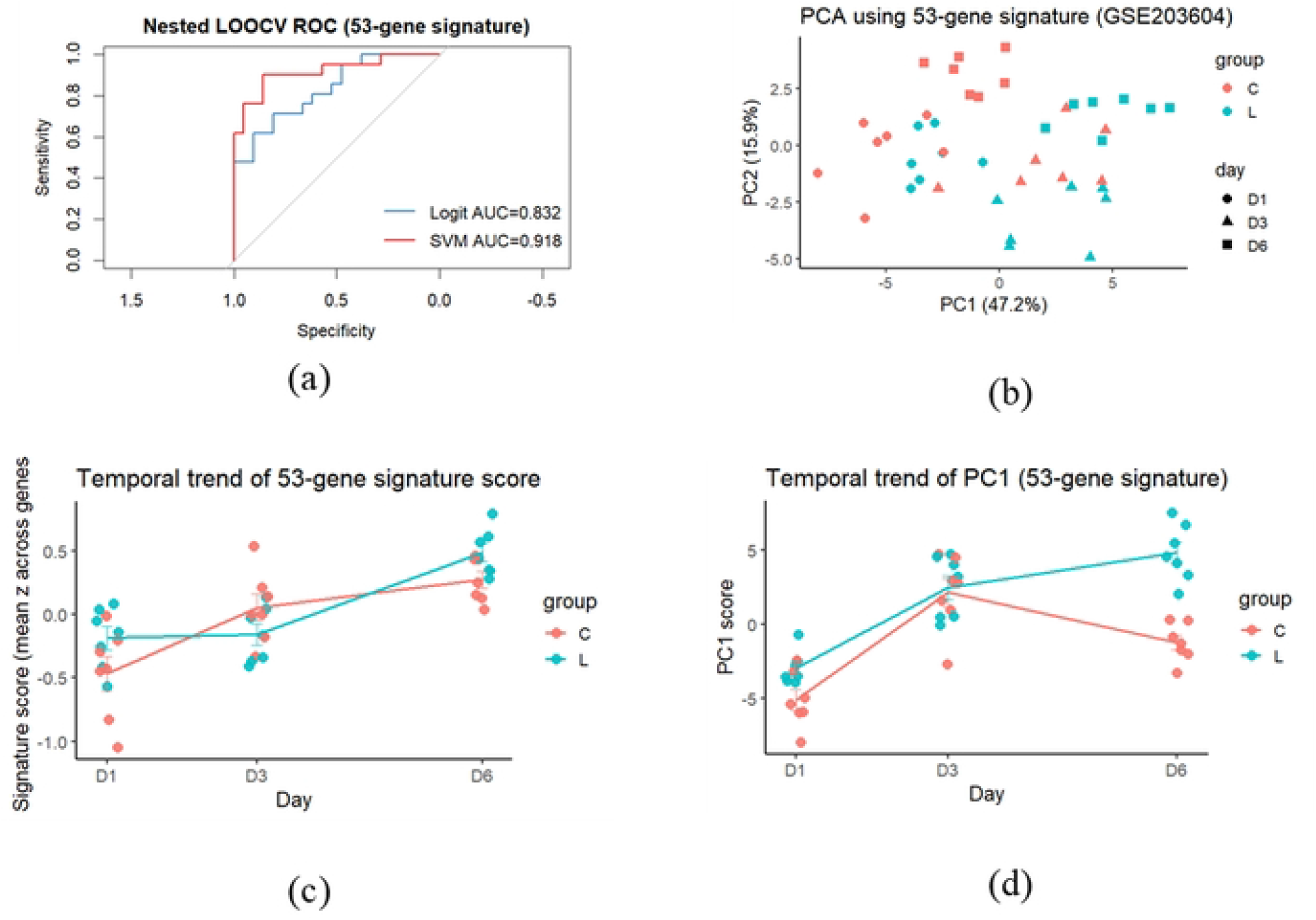

### 3.6 Functional Reorganization of Myopia Gene Programs from Chicken to Human

Gene Ontology enrichment analysis of Boruta-selected genes revealed distinct but related functional profiles between chicken and human orthologs. After redundancy reduction, enriched biological processes in chicken were summarized into parent programs associated with cellular metabolism, ion transport, intracellular trafficking, and post-transcriptional regulation (**Supplementary Data3**). In contrast, human orthologs were enriched for programs related to neuroendocrine regulation, photoreceptor maintenance, hormonal response, and mRNA stability **(Supplementary Data4)**.

Treemap visualization highlighted differences in functional composition between species (see **Supplementary Data5**). Alluvial analysis further showed structured cross-species relationships, with multiple chicken functional programs converging onto fewer higher-order regulatory programs in humans, indicating functional reorganization of myopia-associated gene signatures across species as depicted below in Figure 4.

**Figure 4.**
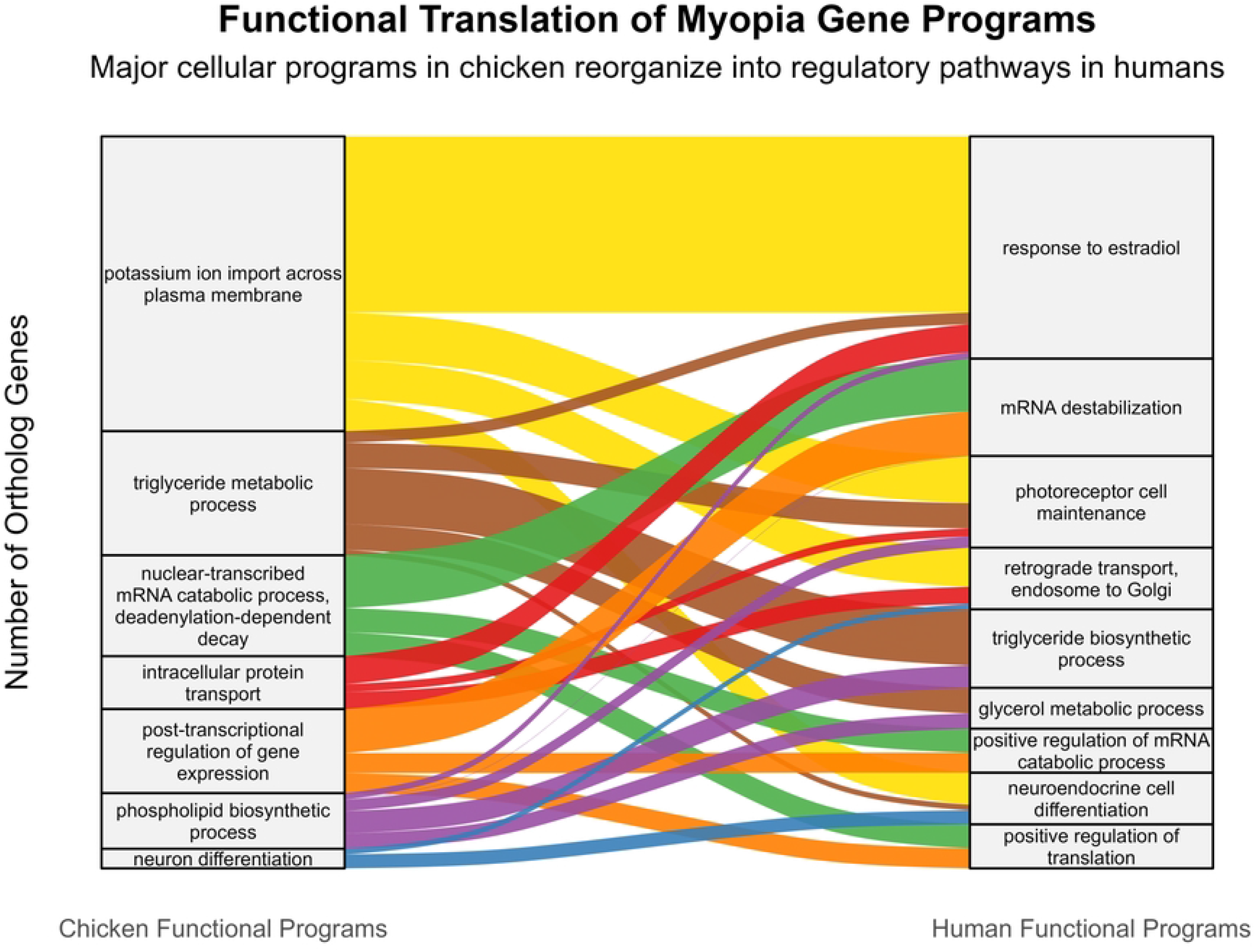

## 4. Discussion

This study identifies ZT8 and ZT12 as circadian-sensitive windows during which myopia-associated transcriptional activity is most pronounced. By integrating retinal and choroidal RNA-seq datasets with multistage ML analysis, we identified gene signatures that consistently distinguish these time points across both myopia onset and progression. These findings extend previous transcriptomic studies demonstrating diurnal regulation in experimental myopia and support a central role for circadian biology in refractive development. Importantly, the consistent patterns observed across both tissues further support the existence of a coordinated retina–choroid axis underlying myopia development.

The identification of ZT8 and ZT12 aligns with known circadian physiology in the eye; ZT8 corresponds to peak light exposure and dopamine signaling, whereas ZT12 marks the transition to darkness and melatonin onset. Dopamine has been shown to inhibit ocular elongation, while melatonin may promote eye growth, indicating opposing circadian influences on refractive development [29]. Experimental studies further demonstrate that disruption of circadian rhythms or light–dark cycles alter myopia progression [30].The strong classification performance observed in this study suggests that these temporal transcriptional differences are both biologically meaningful and physiologically relevant.

Our results also indicate that myopia onset and progression are characterized by distinct molecular programs. Early-stage responses appear to involve rapid retinal signaling and transcriptional regulation, whereas later stages are associated with sustained metabolic and structural remodeling, particularly in the choroid. This transition reinforces the concept of a retina–choroid axis, in which initial retinal responses to visual defocus propagate to downstream tissues, driving structural adaptation of the eye [31]. These findings are consistent with recent RNA-seq studies showing stage-specific transcriptional dynamics during myopia development [14, 15].

Cross-tissue validation demonstrated strong transferability of gene signatures between retina and choroid, providing computational support for shared regulatory mechanisms. Physiologically, the retina acts as a primary circadian oscillator, while the choroid exhibits diurnal changes in thickness and blood flow in response to retinal signaling [32]. The ability of the models to generalize across tissues supports the concept that circadian regulation operates at a systems level, linking retinal activity to choroidal responses within a coordinated retina–choroid axis.

Cross-species analysis (Figure 4) revealed a functional shift in human orthologs, from cellular processes in chickens to higher-order regulatory pathways, including neuroendocrine signaling, photoreceptor maintenance, and hormonal response [33]. These findings suggest that conserved molecular components are embedded within more complex regulatory networks in humans. This functional reorganization highlights species-specific differences while supporting the relevance of the chick model for studying fundamental mechanisms of myopia [13].

An important methodological observation is that ML predictive performance was driven primarily by multivariate expression patterns rather than individual gene effects. The superior performance of RF and SVM models compared to LR, particularly in external validation, indicates that circadian gene programs are encoded through complex, non-linear interactions [34]. This is consistent with broader findings in transcriptomic analyses, where higher-order gene interactions contribute to biological phenotypes[35].

External validation in an independent dataset confirmed that the identified gene signature retains predictive capacity despite differences in experimental design and partial gene overlap. Although only a subset of genes was detected, the signature remained informative, suggesting robustness of the underlying biological signal. The absence of strong univariate differences further supports the interpretation that coordinated gene expression patterns, rather than single-gene effects, drive classification performance [36].

Several limitations should be acknowledged. This study relies on publicly available RNA-seq datasets with relatively modest sample sizes, reflecting the constraints of circadian experimental designs requiring terminal tissue collection. Although the chick model is well established, species-specific differences may limit direct translation to human myopia. In addition, the analysis was restricted to transcriptomic data, and other regulatory layers, including proteomic and metabolic processes, were not assessed. Functional validation experiments were not performed, and circadian sampling was limited to a single 24 h cycle per stage, which may not capture longer-term rhythmic dynamics.

These findings raise the possibility that the efficacy of anti-myopia interventions may be influenced by circadian timing. In particular, if retinal and choroidal signaling is most dynamic between ZT8 and ZT12, aligning atropine administration with this mid-day window may enhance its biological effect. Although this remains speculative, the proposed chronotherapeutic approach warrants further experimental investigation. More broadly, these results highlight the importance of considering time-of-day effects in both experimental design and clinical interpretation. Circadian critical windows may represent periods of increased susceptibility to visual stimuli or enhanced responsiveness to therapeutic interventions. Future studies integrating multi-omics approaches, functional validation, and chronotherapeutic strategies will be essential to translate these findings into clinical applications.

## 5. Conclusion

This study demonstrates that circadian timing is a key determinant of myopia-associated transcriptional activity and provides integrative evidence for a coordinated retina–choroid axis in refractive development. By combining temporal transcriptomics with ML analysis, we identified robust gene signatures that define circadian critical windows (ZT8 and ZT12) and generalize across tissues and disease stages. These findings highlight circadian regulation and retina–choroid interactions as central mechanisms in myopia development and suggest that time-dependent strategies may enhance future approaches to myopia control.

## Supplementary Materials

The following supporting information can be downloaded at:

**Supplementary Data 1:** Phenotypic data.

**Supplementary Data 2:** Genes selected by Boruta feature selection.

**Supplementary Data 3:** Gene Ontology enrichment results for chicken orthologs.

**Supplementary Data 4:** Gene Ontology enrichment results for human orthologs.

**Supplementary Data 5:** TreeMap visualization of functional enrichment results.

## Author Contributions

All authors contributed to the study conception and design. Data analysis and interpretation were primarily performed by Akarapon Watcharapalakorn, under the supervision of Tanakamol Mahawan. Akarapon Watcharapalakorn drafted the manuscript, with substantial revisions and intellectual input from Tanakamol Mahawan. Teera Poyomtip, Patarakorn Tawonkasiwattanakun, Putri Krishna Kumara Dewi, and Thotsapol Thomrongsuwannakij contributed to data interpretation and manuscript review. All authors read and approved the final manuscript.

## Declarations

### Ethics approval and consent to participate

Not applicable.

### Consent for publication

Not applicable.

### Institutional Review Board Statement

This study was reviewed and approved for exemption by the Walailak University Institutional Animal Care and Use Committee (WU-IACUC) under project number WU-ACUC-69005.

### Data Availability Statement

Data Availability Statement: Publicly available GEO datasets **GSE227724, GSE261232**, and **GSE203604** were used in this study. Additional data are provided in the Supplementary Materials. R scripts are available from the corresponding author upon reasonable request.

## Acknowledgments

The authors would like to thank Prof. Richard A. Stone, MD, and his team at the University of Pennsylvania and Prof. Andrei V. Tkatchenko and his team at the Columbia University for generating and publicly sharing the RNA-sequencing data and associated metadata used in this study.

